# New paradigm of CRISPR spacerome for classification of global *Escherichia coli* lineages

**DOI:** 10.64898/2026.01.12.698643

**Authors:** Hanseob Shin, Shri Vishalini Rajaram, Priyanka Singh, Erliang Zeng, David M. Cwiertny, Timothy E. Mattes, Michael A. Pentella, Megan J. Nelson, Ryan T. Jepson, Darrin A. Thompson

**Author notes:** These authors contributed equally to the study as the first authors.

## Abstract

Our research team has recently identified a proof-of-concept for a previously unrecognized spacerome signature in a subset of globally distributed *E. coli*. Motivated by our initial observation, we pursued a more in-depth CRISPR spacerome analysis, which revealed a novel insight in discrimination of global *E. coli* sequence type (ST) lineages. We systematically retrieved publicly available *E. coli* complete genomes, analyzed and compared CRISPR spacerome of globally distributed strains. We found that global strains harbored spacerome that has remained conserved spaceromes for decades across multiple continents. Strains with the conserved spacerome belong to specific ST lineages of clinical importance. In addition, most protospacers were assigned to Gammaproteobacteria and Caudoviricetes. Our findings reveal unexpected long-term conservation of CRISPR spaceromes and their potential as high-resolution markers for *E. coli* epidemiological tracking.

## Introduction

CRISPR is an adaptive immune mechanism found in bacteria and archaea, enabling protection against invading genetic elements such as bacteriophages and plasmids^1^. CRISPR loci are composed of short repetitive DNA sequences interspaced with unique sequences derived from previous foreign DNA encounters, known as spacers. These spacers serve as a genetic memory that allows bacteria to recognize and cleave homologous sequences as an immune system. CRISPR spacerome is defined as the collective repertoire of CRISPR spacers, repeat sequences, and 5’/3’ end flanking regions within a given bacterial species^3^. A previous CRISPR spacerome study found that environmental *Escherichia coli* strains partially shared spacers^4^, while some carbapenemase-producing *E. coli* strains possessed conserved CRISPR spaceromes^3^, which necessitates more comprehensive genomic data analysis by investigating global *E. coli*.

The degree of spacerome conservation or turnover across diverse *E. coli* populations may reveal the dynamics and evolution of CRISPR spacerome. Advances in bioinformatics can fill this knowledge gap, enabling the large-scale characterization of CRISPR spacerome and revealing patterns of conservation and diversification among *E. coli* lineages. Integrating CRISPR spacerome data with phylogenetic genomic analyses will expand our understanding of CRISPR systems.

Tracking clinically significant *E. coli* is crucial for effective infection control. Phenotypic methods, including culture-based screening and susceptibility testing, offer cost-effective options for resource-limited settings^5^. In addition, molecular techniques like whole-genome sequencing (WGS) provide more accurate characteristics and dissemination of carbapenemase producing *E. coli* (CPE)^6^. For example, analysis of single nucleotide polymorphism (SNP) of WGSs has been applied to track the CPE transmission, together with identification of sequence types (STs). There are clinically significant CPE clones, such as ST410, ST131, ST167, and ST405, heralding an emerging high-risk to public health^6,7^. However, current approaches remain insufficient for fully resolving patterns of transmission, calling attention to complement approaches for effective global surveillance^8,9^. Although such studies suggest that targeting a specific STs could be a strategy, they also argue the need to strengthen surveillance approaches to apply higher-resolution methods^8,9^. These findings emphasize the need for novel frameworks that can support conventional approaches.

Recently, the detection of conserved specific CRISPR spacers has been proposed for tracking CPEs and epidemiological studies^3^. This is notable because, although CRISPR spaceromes are theoretically expected to be dynamic and transient due to their acquisition mechanism^10^, some arrays remain conserved across lineages, enabling their use as stable molecular markers^3^. A recent study by Zhang *et al* supports our conclusion, showing that CRISPR spacer acquisition events rarely occur in the human gut environment^11^. In the gut, bacteriophages are mostly lysogenic^12^, which may turn off CRISPR systems in *E. coli*. This repression would reduce the incorporation of new phage-derived spacers and help maintain a stable CRISPR spacerome over time. By contrast, in natural environments where lytic phages are abundant^13^, CRISPR systems are more likely to be activated to defend against infection, promoting spacer acquisition. The reduced spacer acquisition under lysogenic conditions therefore helps preserve existing spacer content, while certain lineages may retain spacers that offer long-term immunity against prevalent phages or plasmids, contributing further to spacerome stability.

In this study, we employed a 3-tier genome mining workflow and found multiple conserved spaceromes persisting for nearly one hundred years in globally distributed *E. coli*. These spaceromes were consistently mapped to clinically important ST lineages, with protospacers identified from the reference sequences of the same plasmids or human gut virome. Our results reveal unexpected conservation of the CRISPR spacerome, which has potential to track *E. coli* at high resolution.

## Results

### Distribution of the CRISPR system and subtype classification across publicly available *E. coli* genomes

From NCBI, where 347,620 genomes have been deposited as of July 11th, 2025, a total of 677 *E. coli* genomes were retrieved with the strict criteria. Among the 677 *E. coli* genomic data, 532 (78.6%) were positive for CRISPR (**Figure 1A**). Of 532 genomes, human-origin *E. coli* genomes were predominant as 53.4% (n = 284), followed by chicken (9.2%, n = 49), pig (8.3%, n = 44), cattle (7.3%, n = 39), dog (3.0%, n = 16), duck (2.8%, n = 15), and minors (16.0%, n = 85) (**Figure 1B**). In addition, among the 532 *E. coli* genomes analyzed, 429 (80.6%) harbored two CRISPR arrays, while 89 genomes (16.7%) contained a single array. Eight (1.5%) and six (1.1%) genomes carried three and four arrays, respectively (**Figure 1C**). The most predominant CRISPR subtype was I-E (n = 446, 90.7%), followed by I-F (n = 31, 6.3%), unidentified (ND, n = 9, 1.8%) and I-E/I-F (n = 6, 1.2%) (**Figure 1D**).

**Figure 1.**
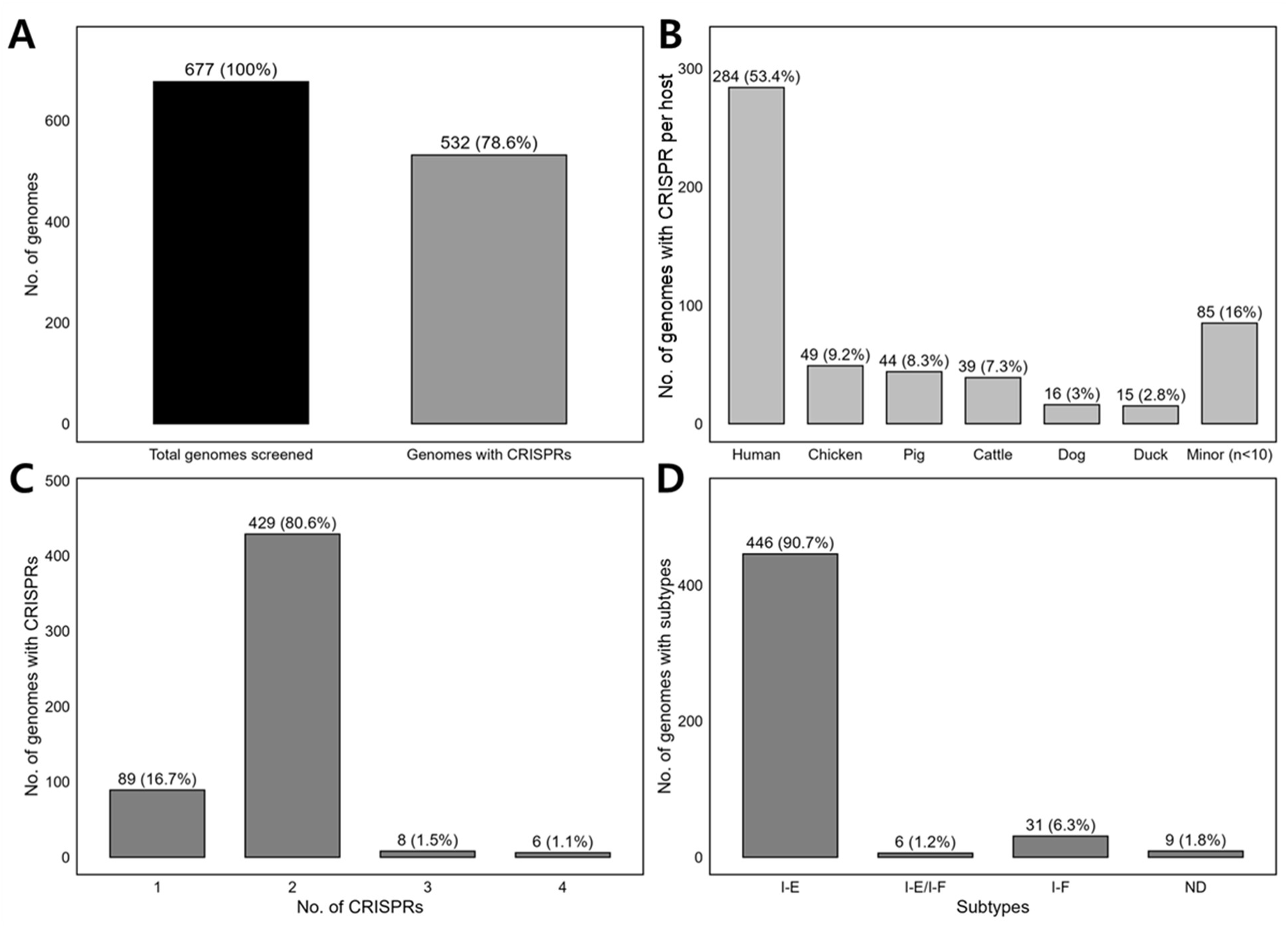
Summary of CRISPR system presence, host origin, the number of CRISPR arrays, and subtype classification across publicly available *E. coli* genomes. (A) Proportion of *E. coli* genomes with and without CRISPR arrays. (B) Distribution of *E. coli* genomes by isolation source (e.g., human, animal, environmental) stratified by presence of CRISPR. (C) The number of CRISPR arrays per *E. coli* genome with at least one array. (D) Classification of CRISPR-Cas subtypes among CRISPR-positive genomes.

### Identification of conserved CRISPR spacerome

The quality of spacers identified was evaluated based on their length and entropy. The distribution shows a strong concentration of spacers around ∼ 32 bp with intermediate entropy values (∼1.85 – 1.95), consistent with the constrained length and compositional diversity characteristic of *E. coli* CRISPR spacers at the population level **(Figure S1).** Phylogenetic reconstruction of 532 WGS and 997 CRISPR sequences annotated with host and geographic metadata revealed that several CRISPR groups appeared to form clusters (**Figure S2 and S3**). While rough cluster construction suggested potential conservation across lineages, the overall sequence divergence was substantial, and thus, conserved spacerome could not be explicitly discerned. This observation motivated a more stringent comparison, driving the application of Jaccard network analysis to identify the conserved CRISPR spacerome. Network analysis of shared spaceromes identified the conserved CRISPR spaceromes in 20 *E. coli* groups (**Figure 2A**). Specifically, 10 groups (A – J) were selected based on their relevance to public health, representing human – human or human – livestock animals. Within each group, *E. coli* strains with conserved CRISPR spaceromes were identified across multiple countries, years of isolation, and host sources.

**Figure 2.**
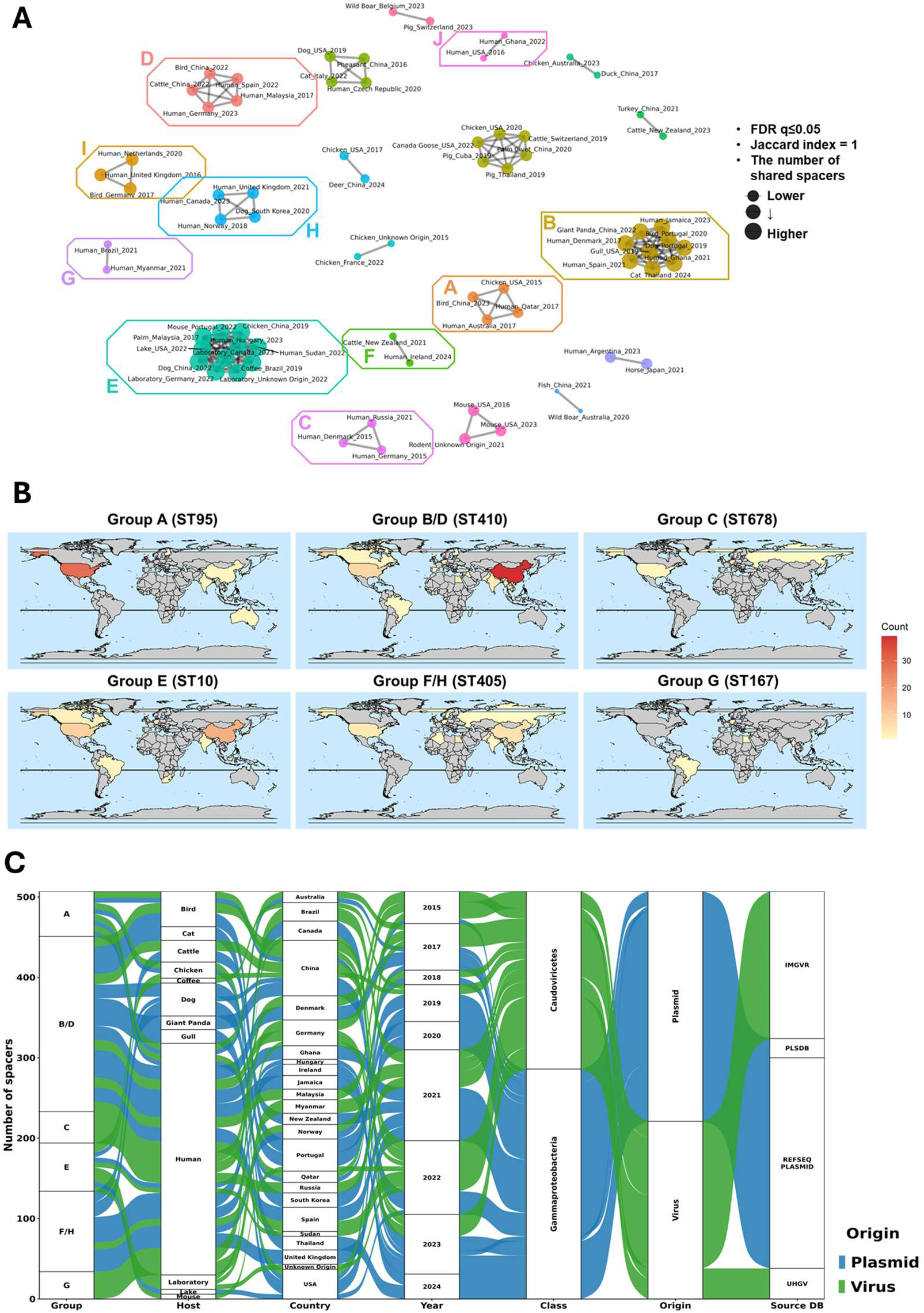
CRISPR Spacerome-Based Clustering and Global Distribution of *E. coli* Lineages. (A) Conserved CRISPR spaceromes were identified using the Jaccard similarity index. Pairwise connections were retained only when the FDR-adjusted *P*-value was ≤ 0.05 and the Jaccard index equaled 1. Each node represents an isolate’s CRISPR spacerome, and edges indicate complete spacerome identity (Jaccard index = 1). Node size reflects the number of shared spacers, while node color denotes one of the six major spacerome groups (A – G) selected from 20 total groups based on relevance to human health. Connected nodes represent isolates that share conserved CRISPR spaceromes. (B) Global distribution of 475 *E. coli* strains harboring conserved CRISPR spaceromes between 1922 and 2024. Six major groups (A–G) identified by spacerome analysis are shown. Frequency of each group per country is represented by color intensity, with red indicating high frequency and light yellow low frequency. Countries with no detected strains are shown in grey. (C) Mapping of CRISPR spacer origins (protospacers) across *E. coli* groups. Each spacer is traced from its host category (Host) across various countries (Country) and year (Year) through the taxonomic class of the matched protospacer (Class) to the predicted genetic origin element (Origin), and to the reference database (Source DB). Green and blue flows represent spacers mapped to viral- and plasmid-derived sequences, respectively.

For example, in group A, *E. coli* strains with a conserved CRISPR spacerome were identified in a chicken from the USA (2015), humans from Australia and Qatar (2017), and a bird from China (2023). Elsewhere, in group C, another distinct conserved spacerome was detected in two human-derived strains from Russia (2021) and Denmark (2015). Likewise, in the other groups, different conserved spaceromes were identified between humans, or humans and livestock animals. Groups B and D, as well as groups H and F, were initially classified as distinct based on differences in their CRISPR spacerome sequences. Comparative analysis revealed that groups B and D, and H and F differed only in the 5′ flanking region sequences, in which group B and F has shorter 5’-end flanking region sequences. Despite these differences in the 5’-end flanking regions sequences, whole spacerome BLAST search consistently retrieved the same strains from both groups. Therefore, groups B – D and F – H were consolidated. The detailed spacerome sequences are provided in **Table S1**. Finally, eight different conserved spaceromes (of ten groups) were determined for downstream analysis as follows: A, B/D, C, E, F/H, G, I, J.

### Genomic and CRISPR spacerome analysis of expanded global *E. coli*

Across the eight groups, several core *cas* genes were consistently identified **(Table 1)**. Variability was observed in the subtype-specific signature genes: *csy1*–*3* were unique to group A (type I-F), whereas *cse1* and *cse2* were present in groups C, E, F/H, G, I and J (type I-E). While the majority of groups (type I-E) harbored a standardized gene organization, group A was distinguished by the presence of the I-F-specific *csy* gene set.

**Table 1.** Classification of *E. coli* genomes based on conserved CRISPR spacerome groups and sequence types (STs). *E. coli* genomes sharing conserved CRISPR spacerome profiles were grouped (A – J) and summarized by the number of matching genomes, dominant multilocus sequence types (major STs), and less frequent lineages (minor STs). For each group, the associated CRISPR–Cas subtype (I-E or I-F), Cas protein–coding gene composition, and the number of spacers detected in each CRISPR array are shown. Spacer counts refer to the representative CRISPR arrays conserved within each spacerome group.

| Group | Genomes matching spacerome | Major ST | Minor STs | Type | Cas protein coding genes | CRISPR array 1 | CRISPR array 2 |
| --- | --- | --- | --- | --- | --- | --- | --- |
| A | 161 | ST95 (n = 145) | ST2619 and unidentified (n = 5), ST416 (n = 3), ST1231, ST11777, ST643 (n = 1) | I-F | *cas1, cas3, cas2, csy1, csy2, csy3, cas6* | CRISPR array (8 spacers) | CRISPR array (6 spacers) |
| B/D | 115 | ST410 (n = 93) | ST23 (n = 13), ST2851 (n = 4), unidentified ST (n = 3), ST1998 and ST9862 (n = 1) | I-E | *cas3, cas7, cas5, cas6, cas1, cas2* | CRISPR array (8 spacers) | CRISPR array (9 spacers) |
| C | 30 | ST678 (n = 24) | ST602 (n = 6) | I-E | *cas3, cse1, cse2, cas7, cas5, cas6, cas1, cas2* | CRISPR array (13 spacers) | — |
| E | 108 | ST10 (n = 87) | ST1060 (n = 14), unidentified ST (n = 6), ST11066 (n = 1) | I-E | *cas2, cas1, cas6, cas5, cas7, cse2, cse1, cas3* | CRISPR array (12 spacers) | CRISPR array (6 spacers) |
| F/H | 50 | ST405 (n = 40) | Unidentified ST (n = 4), ST617 (n = 3), ST648, ST1284, ST4204 (n = 1) | I-E | *cas3, cse1, cse2, cas7, cas5, cas6, cas1, cas2* | CRISPR array (5 spacers) | CRISPR array (9 spacers) |
| G | 11 | ST167 (n = 7) | ST8453 (n = 3), unidentified ST (n = 1) | I-E | *cas3, cse1, cse2, cas7, cas5, cas6, cas1, cas2* | CRISPR array (6 spacers) | CRISPR array (11 spacers) |
| I | 2 | ST648<br>(n = 2) | - | I-E | *cas3, cse1,*<br>*cse2, cas7,*<br>*cas5, cas6,*<br>*cas1, cas2* | CRISPR array (6<br>spacers) | CRISPR array (11<br>spacers) |
| J | 4 | ST617<br>(n = 4) | - | I-E | *cas3, cse1,*<br>*cse2, cas7,*<br>*cas5, cas6,*<br>*cas1, cas2* | CRISPR array (23<br>spacers) | CRISPR array (13<br>spacers) |

A subsequent BLAST search was performed using the complete spacerome sequences as queries, which identified a total of 1,746 WGSs matching spacerome sequences from the six groups. After removal of redundant hits and filtering for genomes carrying two spacer arrays (if two are present) to secure only the overlapping WGSs, we obtained a final dataset: group A (n = 161), group B/D (n = 115), group C (n = 30), group E (n = 108), group F/H (n = 50), group G (n = 11), group I (n = 2), and group J (n = 4) resulting in 481 unique WGSs in total (**Table 1**). Of eight groups, only six (A, B/D, C, E, F/H and G) were subject to further analysis since the number of genomes matching spacerome in group I and J were too low as two and four.

### Global distribution of CRISPR spacerome-defined *E. coli* lineages

Phylogenetic analysis based on multi-locus sequence type (MLST) demonstrates that the majority of isolates in each CRISPR spacerome group belonged to specific STs, namely ST95 in group A (n = 145/161), ST410 in group B/D (n = 93/115), ST678 in group C (n = 24/30), ST10 in group E (n = 87/108), ST405 in group F/H (n = 40/50), and ST167 in group G (n = 7/11) (**Table 1**). Minor STs were also detected within each group. The global survey of *E. coli* lineages from 1922 to 2024, conveyed by CRISPR spacerome, showed geographic distribution (**Figure 2B**). Overall, the higher occurrence frequency was shown in North America, Europe, and East Asia. In contrast, relatively few lineages were detected in Africa and South America. In addition, the temporal occurrence of a total of 475 *E. coli* strains in six groups (A, B/D, C, E, F/H, and G) was visualized in **Video 1**. These lineages were broadly distributed across continents and isolated over nearly a century, from 1922 to 2024. The detailed information about strain ID, metadata (year and country of isolation), and accession numbers is provided in **Table S2**.

Across the conserved CRISPR spacerome groups, distinct but overlapping global dissemination patterns were observed (**Table S2**). Group A (ST95-related) emerged in the United States in the 1970s and later showed repeated expansion in the United Kingdom during the 2010s, followed by spread into Northern Europe and parts of Asia. Group B/D (ST410-related) originated early in Europe and West Africa, expanded rapidly into Asia after 2010, and subsequently disseminated broadly across Europe, the Americas, and Africa, representing one of the most globally distributed lineages. Group C (ST678-related) appeared in the early 2000s in Germany and South Korea and has since shown intermittent but persistent circulation mainly across Europe and East Asia. Group E (ST10-related) represents the oldest lineage, with reports dating back to the early 20th century in North America, later expanding into Asia and subsequently re-emerging across Europe, the Americas, and Africa. Group F/H (ST405-related) was first detected in Northern Europe and has undergone extensive expansion over the past decade across Asia, Europe, Africa, and North America. Group G (ST167-related) is the most recently emerged lineage, first detected in West Africa in 2016, yet it has already achieved rapid intercontinental spread across Africa, Asia, Europe, and South America within a short time frame.

### Comparison of CRISPR spacerome within identical ST lineages

The spaceromes were compared among isolates belonging to the same ST lineages in order to evaluate whether the identical CRISPR spaceromes observed across isolates were influenced by lineage-specific bias (**Table 2**). Within each ST, several isolates displayed identical or nearly identical spacer arrays, whereas others showed partially shared or completely distinct patterns.

**Table 2.**
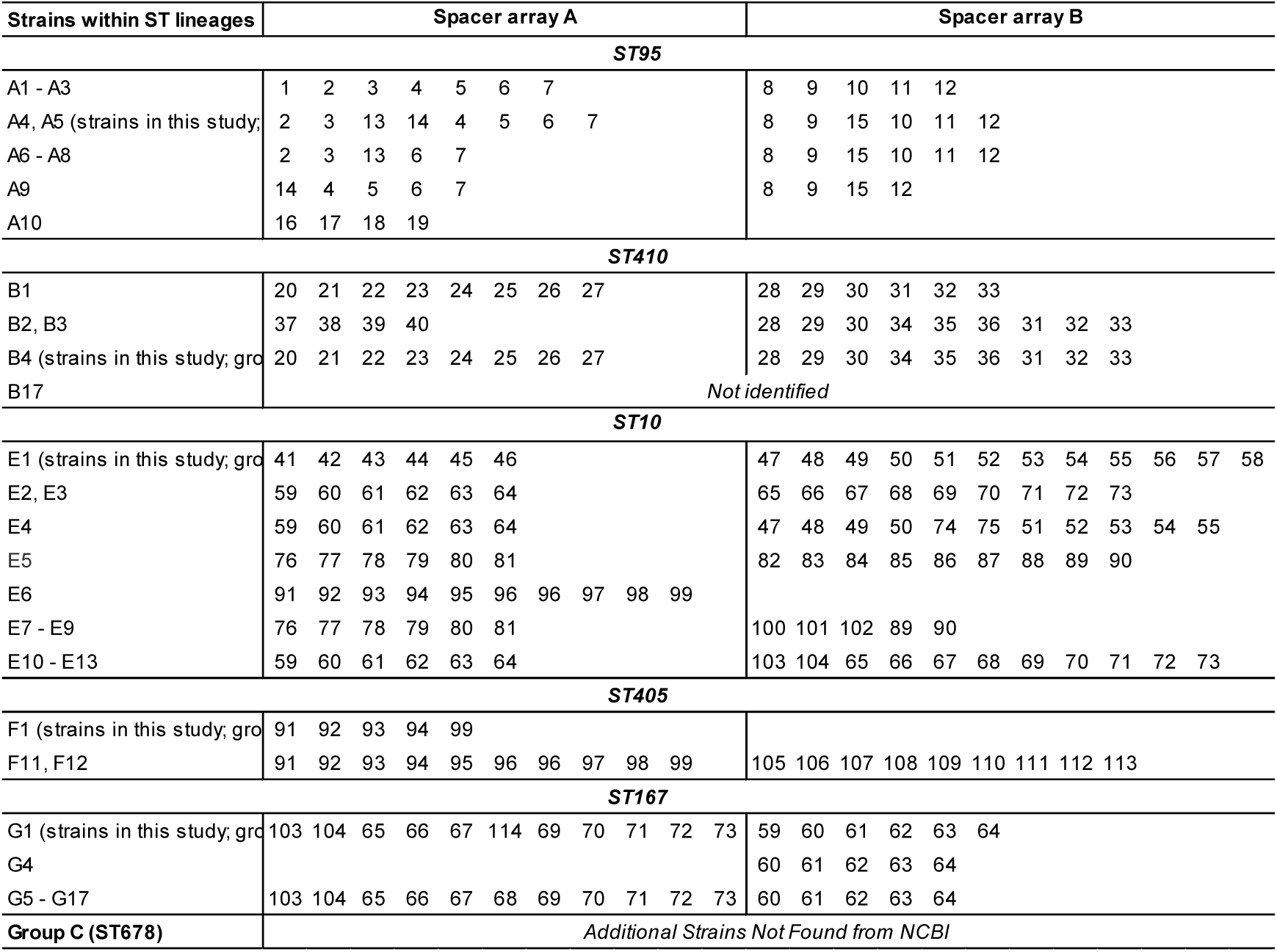
Spacer array structures across major E. coli sequence type (ST) lineages. The CRISPR spaceromes within each group are summarized. For each ST lineage, strains are classified by spacerome-defined sublineages, and spacer arrays are separated into array A and B when present. Spacer numbers correspond to unique spacer IDs assigned in this study. CRISPR arrays were detected using CRISPRFinder with default parameters. When multiple CRISPR arrays occurred in a genome, they were distinguished by suffixes (e.g., A1 – A3, E1 – E13). All repeat and spacer sequences are shown in the 5′ → 3′ orientation. Strains listed as Not identified indicate that no CRISPR array was detected. Group C (ST678) includes additional strains not available from NCBI.

Within ST95 lineage, isolates A1 – A3 shared an identical spacerome (spacer 1 – 7 and 8 – 12), while A10 exhibited one different array (spacers 16 – 19). Other isolates (A4 – A9) possessed partially identical spacers. Within ST410, identical spacers (spacers 20 – 27 and 28 – 33) were commonly identified in multiple isolates (B1, B4 – B16), with an extended array (spacers 34 – 36) detected in some strains (B4 – B16). Two strains (B2 and B3) within ST410 also possessed the identical spacer array B with B4 – B16 while spacer array A was completely different, harboring spacer 37 – 40. In ST10, both identical and divergent arrays were present.

While our strains (designated as E1) in this study carried spacers 41 – 46 and 47 – 58, isolates E6 contained only one extended array (spacers 91 – 99). The other isolates within ST10 lineage also carried distinct spacer compositions. ST405 strains carried two different arrays (F1, F2 – F10: 91 −94, 99, and F11, F12: 91 – 99 and 105 – 113). All ST167 isolates commonly carried spacers 60 −64 while the other isolates (G1 – G3 and G5 – G17) also possessed. The ST167 lineage exhibits two major structural patterns, with G1 – G3 maintaining a full-length conserved array composed of spacers 103 – 104, 65 – 67, 114, 69 – 73, and 59 – 64. Several spacers, including spacers 65–73, were shared across ST10 and ST167 lineages. In contrast, some isolates showed no detectable CRISPR array, and the CRISPR locus was not found in ST678 genomes retrieved from NCBI.

### Protospacer identification

Protospacers corresponding to *E. coli* CRISPR spacers from six CRISPR groups (A – G) were resolved across taxonomic class, molecular origin, host category, and reference database (**Figure 2C**). Most spacers mapped to two dominant Class-Origin categories, Caudoviricetes-virus and Gammaproteobacteria-plasmid, whereas a small fraction showed support for both categories or lacked identifiable protospacers. Protospacer origin was strongly structured by CRISPR group. Groups C and G were predominantly enriched for viral protospacers dominated by Caudoviricetes, whereas the other groups mixed viral and plasmid origin profiles. Host-associated patterns were secondary to group-level structure. Non-human host–associated isolates, driven primarily by bird-derived samples, were predominantly associated with Caudoviricetes-derived protospacers. In contrast, isolates from other non-human hosts were enriched for plasmid-associated protospacers. Human-associated isolates contributed substantially to both viral and plasmid categories, resulting in the most heterogeneous origin profiles observed across host categories. Protospacer assignments were supported primarily by IMG/VR for viral matches and RefSeq Plasmid for plasmid matches, with smaller contributions from PLSDB and UHGV.

## Discussion

Building on our prior findings suggesting that identical spaceromes may enable tracking of global *E. coli* dissemination^3^, we tested this concept by analyzing a large collection of global *E. coli* genomes and their spaceromes worldwide. Our CRISPR screening revealed that most *E. coli* genomes in NCBI database harbor CRISPR arrays. In contrast, previous studies described that only approximately 10% of microorganisms possess CRISPR-Cas systems, in which these analyses primarily focused on 1,724 microorganisms derived from metagenome-assembled genomes in the NCBI database, rather than *E. coli* specifically^14^. Another investigation found that 4 and 38% of *E. coli* strains from human and animal feces carried CRISPR loci, respectively^4^. However, those prior studies investigated strains regardless of clinical significance, whereas all CRISPR-positive strains analyzed in this study were obtained from NCBI, where the majority of deposited genomes are clinically relevant. It is important to note that *E. coli* genomes deposited in NCBI are largely biased towards human-associated isolates selected for their clinical relevance. This sampling bias may influence the observed frequency of CRISPR array because genomes isolated from natural environments are less likely to be clinically significant and therefore might be underrepresented in NCBI database. Hence, our results may overestimate CRISPR-array prevalence in clinically relevant strains relative to *E. coli* populations that are not routinely deposited to in publicly available genome resources.

Interestingly, among the strains positive for CRISPR system, most of them possess two CRISPR arrays, with few strains harboring up to four arrays. Recent large-scale comparative analyses suggested that multiple CRISPR arrays frequently coexist within a single bacterial genome^15^. Despite such high prevalence, the justification of multiplexity has not been resolved. It has been only assumed that multiple arrays represent a strategy to maximize efficiency of CRISPR-Cas system by optimizing the spacer length and the number of arrays^16^. Theoretically, longer CRISPR arrays allow the bacterial cell to preserve older spacers, which results in extending the immune memory^16^. However, too long arrays may reduce the efficiency of expressing new spacers due to the distant location from the associated *Cas* proteins, which can slower the response to external attack. Conversely, shorter arrays tend to lose older spacers rapidly but enable faster acquisition and deployment of new spacers^16^. This way, multiple different arrays within a genome may complement each other, with two modules: long memory with slow-learning and short memory with fast-learning. Moreover, a prior comparative genomic evidence showed that CRISPR spacer arrays are positioned on both sides of the *Cas* locus, and the length of arrays is mostly limited to 50 spacers^15^. Taken together, although these explanations remain hypothetical, these findings support the multiplexity of CRISPR arrays.

Consistently with those prior studies, our spacerome analyses also identified the coexistence of two CRISPR arrays in global *E. coli* genomes with different lengths. However, both arrays in our *E. coli* collection remained identical for decades in multiple countries, which appears inconsistent with the expectation of a dynamic, fast-learning module. This observation drove a following hypothesis: “*Given that most of the conserved arrays were found in E. coli of human or animal gut environments, it is plausible that the intestinal environment, characterized by abundant lysogenic phages and predominantly commensal microbial communities, reduces selective pressure for inactivation of spacer acquisition or loss*”. Under these conditions, CRISPR arrays may become inactivated and stably maintained within the gut bacterial genome, resulting in the long-term conservation that we observed in this study.

An interesting pattern was also found in the distribution of CRISPR subtypes (I-E and I-F), among which I-E was found to be predominant in our *E. coli* collection. While subtype I-E has not been directly implicated in multidrug resistance in prior studies, subtype I-F is more frequently detected among antimicrobial-susceptible strains and shows a negative association with the acquisition of resistance plasmids, particularly those of the IncF/I incompatibility group^17^. The dominance of I-E subtype in public repositories (NCBI) likely reflects a sampling bias, as researchers preferentially deposit clinically significant isolates, especially those harboring antimicrobial resistance or virulence. Thus, if more comprehensive genomic analysis of *E. coli* from diverse natural environmental metrices may show different distribution patterns of CRISPR subtypes. Then, it should be able to reveal the potential role of *E. coli* CRISPR subtypes in dissemination of antimicrobial resistance or virulence.

Followed by CRISPR spacerome analysis of expanded *E. coli* genome collections, we discovered worthwhile point to discuss, which is that the six conserved CRISPR spaceromes are associated with specific *E. coli* ST lineages. As discussed above, CRISPR spaceromes are generally regarded as dynamic and transient, reflecting the continuous arm race between bacteria and invading phages or plasmids.^10^ However, a recent study suggested CRISPR spacer acquisition is a rare event in the human gut environment^11^, which is an evidence on our novel finding that CRISPR spacerome in our *E. coli* collection has been conserved over decades across continents. Also, conserved spacerome was associated with specific ST lineages.

However, this interpretation could lead to a bias, as it may imply that all isolates within the same ST harbor identical spaceromes although variation or distinct spacerome can still be observed within the same ST lineages. To resolve this bias, we examined the variation of CRISPR spacerome within certain ST lineages. Several isolates within the same ST shared identical, partially identical or completely different spacer composition. For instance, strains within ST95 and ST410 lineages contained highly conserved core spacers, whereas those within ST10 displayed multiple sublineages with distinct spaceromes. Some strains within ST167 had only one different spacer on the arrays with those within ST10. Interestingly, although ST167 and ST10 represent distinct lineages within the ST10 clonal complex, we observed that several ST167 strains shared partially identical spacer contents with ST10 strains. Such occurrence patterns of spacerome within the same ST lineages do not necessarily imply they are clonally related. However, their retention of spacers might be inherited from a common ancestor. The important thing is, that these lineage-specific spaceromes have persisted for decades across globally distributed isolates. This observation supports our conclusion that the conserved spacerome functions as a powerful strain-level discrimination tool, even within a single ST.

The six lineages, harboring conserved spaceromes, are clinically relevant clones. *E. coli* ST95 is a globally disseminated extraintestinal pathogenic *E. coli* lineage responsible for urinary tract infections, bloodstream infections, neonatal meningitis in humans^18,19^. *E. coli* ST410 is an emerging multidrug-resistant clone of global concern that has evolved through distinct clades and subclades^20,21^. ST405 strains are also notable for carrying extended-spectrum β-lactamase genes, and have been associated with clonal dissemination alongside ST131 strains^22^. ST678, closely related to the O104:H4 outbreak lineage, represents another pathogenic clade with epidemiological significance^23^. Finally, ST167 belonging to the clonal complex of ST10 is a rising carbapenemase-producing clone that has been implicated in hospital outbreaks across multiple continents^24^. The worldwide occurrence of these STs heralds global public health, which drives the need for higher resolution of tracking *E. coli* transmission. In brief, conserved CRISPR spaceromes were found to closely align with specific *E. coli* ST lineages, showing their potential as precise markers for tracking *E. coli*. Importantly, not all STs carried the conserved spacerome (identified in this study), which benefits the spacerome analysis. This is because it can help to track at more fine-scale within the same ST, when a conserved array is identified in the further studies.

Notably the protospacer mapping also exposed an additional pattern consistent across the groups. Although CRISPR arrays can theoretically capture sequences from a broad diversity of phages and plasmids^25^, many arrays contained spacers aligning to certain classes (Gammaproteobacteria plasmids and Caudoviricetes). This repetitive alignment pattern suggests that a restricted subset of genetic elements contributed disproportionately to spacer acquisition. Such patterns could arise from recurrent encounters with the same phage or plasmid populations or from localized regions enriched in PAM motifs that function as protospacer hotspots^26^. Recent work has reported a preferential recovery of protospacers from prophage regions^27,28^, although the drivers of this pattern remain unknown. In natural environments, CRISPR-Cas adaptation is expected to sample a wide range of invaders^4^, whereas the CRISPR-Cas system is frequently inactive in the gut ecosystem^11^. The intestinal environment is dominated by lysogenic phages and stable commensal bacterial populations^29,30,31,32^, conditions that collectively impose lower selective pressure for rapid spacer incorporation. Consistently, gut-associated CRISPR-Cas systems in *E. coli* are frequently reported to be inactivated or transcriptionally downregulated, reducing the energetic cost of maintaining active immunity^33,34^. Under such conditions, spacer arrays may persist as conserved genomic features rather than dynamic immune records, becoming stably inherited across lineages over long evolutionary timescales. The predominance of conserved spaceromes among human and animal gut *E. coli* strains observed in our dataset is compatible with this scenario, especially given the relatively stable gut virome and plasmidome compared with more variable external environments^35,36^. Together, these observations support the idea that gut-associated CRISPR spaceromes reflect long-term retention of ancestral arrays, rather than ongoing diversification driven by active immune acquisition.

This study has limitation, which is challenging to overcome, that our genomic analysis depends on public repositories such as NCBI, which are biased toward clinically significant isolates^37^. In contrast, less environmental isolates might be deposited in NCBI. Moreover, bacterial WGSs and deposition into publicly available genomic database from low-income countries might be relatively less active, probably due to the high cost of high-throughput bacterial WGS. The other limitation of comparative spacerome analysis within the same ST lineages is that retrieving all genomes belonging to a given ST remains inherently challenging, as many publicly deposited assemblies lack ST annotations and therefore escape systematic searches in NCBI. Despite these limitations, the consistent detection of conserved arrays for multiple decades across continents supports the robust potential of spacerome analysis. It is noteworthy that spacerome analysis represents a powerful framework by which future studies can extend to map pathogen transmission and resolve fine-scale clonal spread.

Collectively, our findings contribute to establishing a new paradigm of CRISPR spaceromes including what CRISPR spaceromes are and how they are interpreted and applied, at least in *E. coli*. Rather than being transient, certain spaceromes can remain stably conserved within high-risk clonal lineages in the gut environments. This observation shows an advanced knowledge in microbial genetics, particularly for a practical tool for epidemiology. Although numerous global surveillance studies have been conducted to date^7,9,24,38,39^, they have not directly revealed the specific transmission routes for the worldwide spread of specific *E. coli* ST lineages. We identified six conserved spacerome types selected regarding public health and consistently detected across multiple continents for decades. The critical point is not simply the number of genomes investigated, but rather the persistence and recurrence of these conserved spaceromes in diverse geographic and temporal contexts. While our analysis determined only six spaceromes, further characterization of spaceromes from newly isolated strains will help predict the clonal spread of *E. coli* infections, as a high-resolution epidemiological approach.

Thus, we suggest a further direction framework **(see Figure 3)** for applying spacerome analysis to high-resolution epidemiology. First, isolates can be systematically screened for CRISPR arrays through various tools^40–42^, followed by extraction and BLAST search of spacerome sequences. By integrating NCBI BLAST search results with downloaded WGS data and associated metadata, MLST and related analyses would help achieve more effective epidemiological comparisons across regions and years. In addition, for further research, we will build databases of conserved spaceromes to curate *spacerome typing,* linked to high-risk *E. coli* lineages so that researchers can easily identify if their strains are closely linked with globally significant threatening clones. By integrating spacerome analysis into genomic surveillance pipelines, we anticipate a new resolution in tracking the emergence and spread of clinically significant *E. coli* clones.

**Figure 3.**
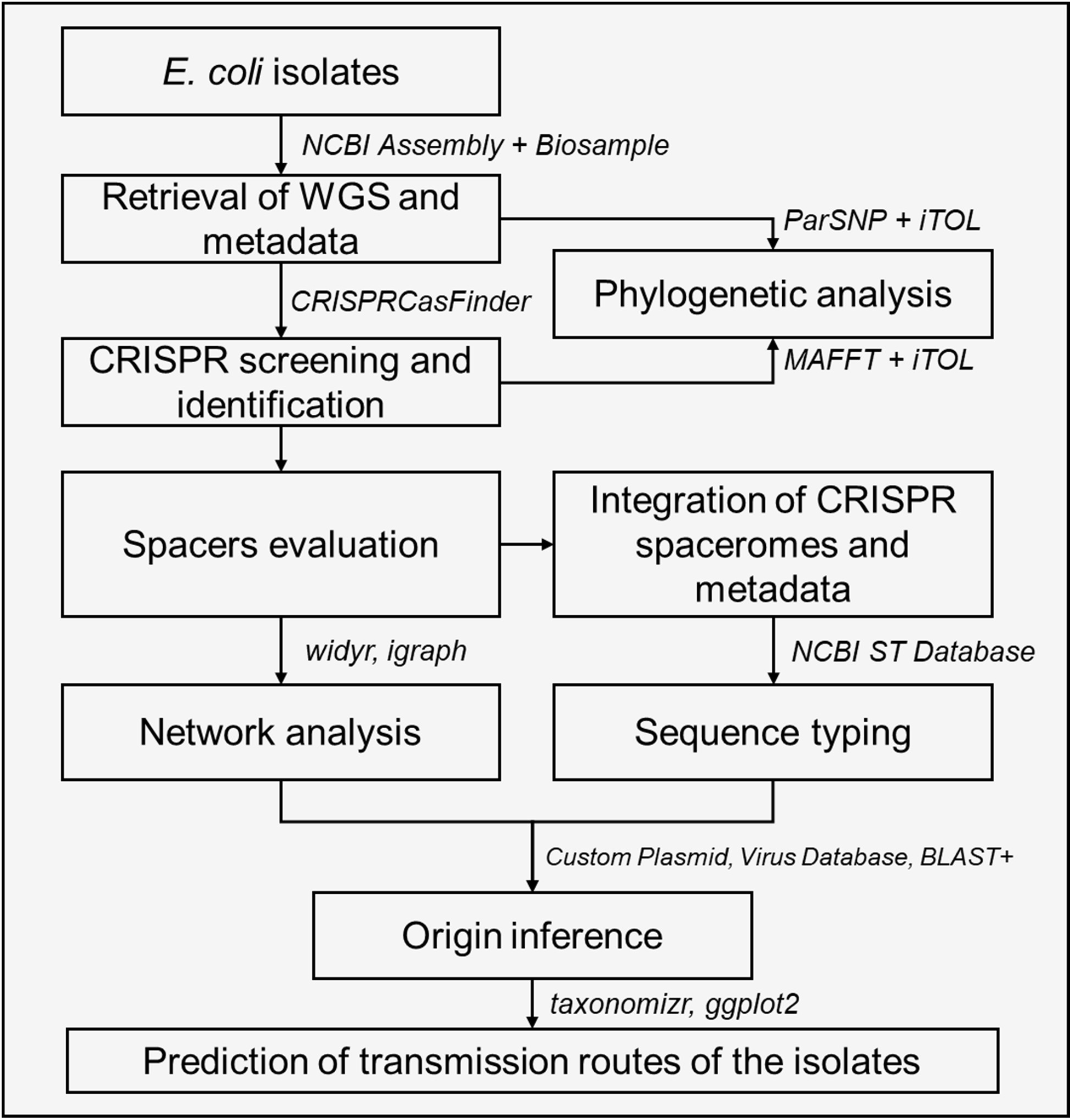
**Framework for CRISPR spacerome-based Genomic surveillance of *E. coli.*** This framework outlines an integrated approach to predict the transmission routes of *E. coli* using CRISPR spacerome analysis. By integrating conserved spacerome sequences with spatiotemporal mapping, genomic analyses, this workflow enables high-resolution tracking of *E. coli* lineages and identification of their potential transmission routes.

## Materials and Methods

### Study design

This study was designed to systematically identify conserved CRISPR spacerome structures across *E. coli* strains and to infer the potential origins and evolutionary routes of these lineages. The workflow consisted of three analytical tiers, together with a comparative spacerome analysis integrating genomic and metadata sources **(Figure 4)**.

**Figure 4.**
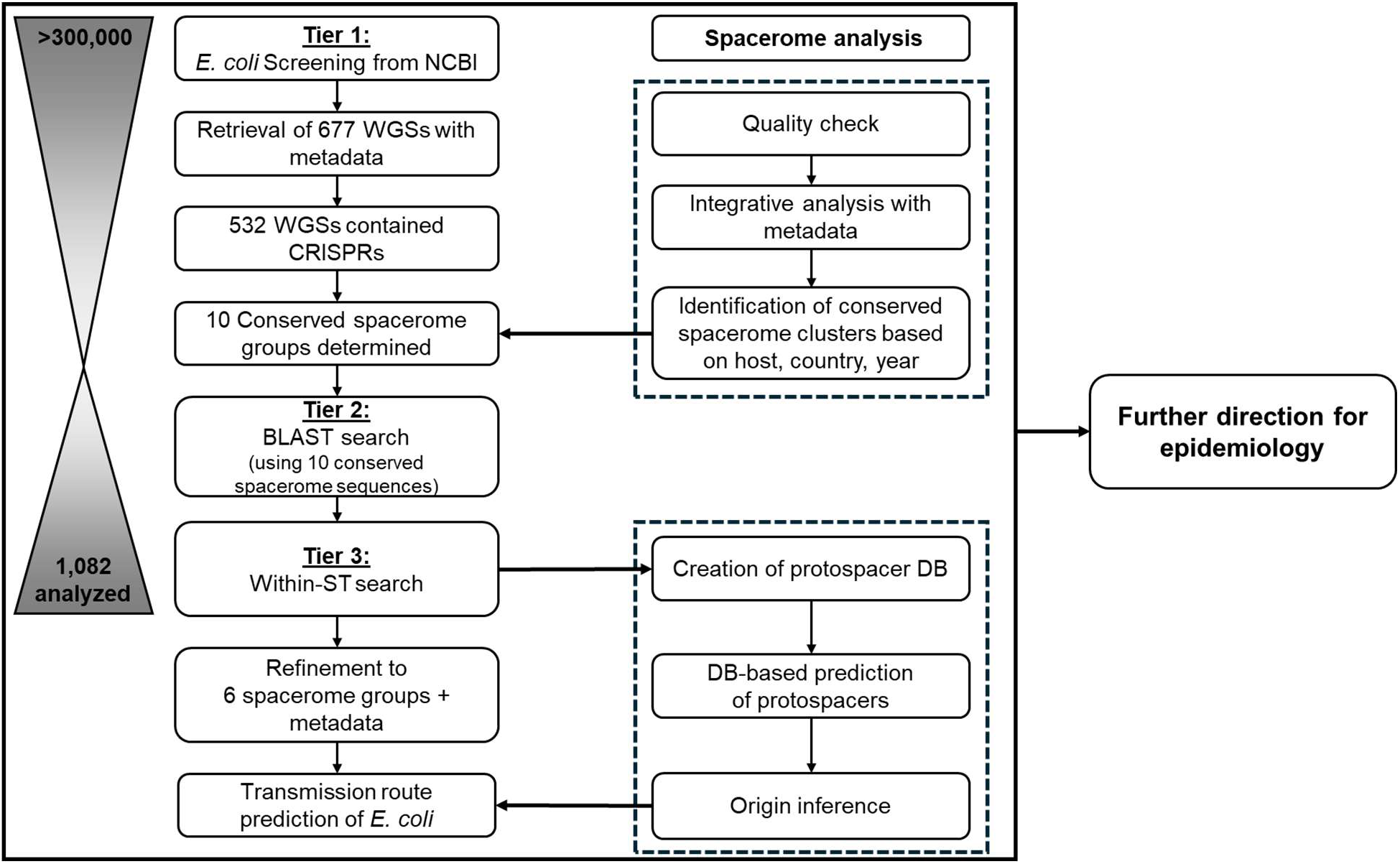
Workflow of CRISPR spacerome analysis in global *E. coli* genomes. Stepwise selection and grouping of *E. coli* whole-genome sequences (WGSs) from >300,000 deposited genomes, including filtering by CRISPR-Cas presence, Jaccard index, and clinical relevance, resulting in eight representative groups. Downstream CRISPR spacerome analysis across isolates of each group, integrating metadata to predict potential transmission routes and future applications, together with protospacer identification, database (DB)-based classification, origin inference, and hypothesis of functional roles.

In Tier 1, we aimed to select *E. coli* genomes that contained analyzable CRISPR spaceromes. Because analyzing the full set of > 300,000 deposited *E. coli* genomes would introduce unnecessary complexity, strict quality criteria were applied to retrieve a manageable, high-confidence dataset for initial spacerome profiling (described below). In Tier 2, the spaceromes identified in Tier 1 were used as query sequences for a broader BLAST-based survey across public *E. coli* genomes. This step was designed to refine the number of conserved spacerome groups. In Tier 3, within-ST spacerome refinement was performed. To resolve fine-scale variation and detect lineage-specific signatures, we conducted the targeted BLAST searches within individual sequence types that matched the conserved spacerome groups. Following the three-tier workflow, we carried out a comparative spacerome analysis integrating genomic metadata, protospacer predictions, and origin inference, followed by further directions and studies.

### Retrieval of genomes

A search in the NCBI Assembly database (https://www.ncbi.nlm.nih.gov/assembly) was conducted using the term “*Escherichia coli*” with filters applied to select only genomes listed under the latest RefSeq version, with complete genome assembly level, RefSeq annotations, and no anomalous or unverified entries. Assembly metadata were downloaded from NCBI in the form of RefSeq assembly reports and parsed to extract relevant information for each genome, including accession numbers, strain names, and assembly statistics.

BioSample accession IDs were extracted from the assembly metadata and submitted to the NCBI Batch Entrez (https://www.ncbi.nlm.nih.gov/sites/batchentrez) to retrieve detailed sample-level metadata, including host, isolation source, geographic origin, and year of collection. These metadata were processed to generate structured annotation tables. In addition to public genomes, a set of *E. coli* in-house strains (“Shin strains”) was manually mapped to BioSample accessions. The corresponding assembly and BioSample records were retrieved and processed using the same approach as for the public RefSeq genomes. Metadata from both sources, RefSeq and Shin, were merged and harmonized into a unified dataset.

### Data scrubbing of retrieved genomes

During curation, entries lacking valid host information, or those with ambiguous or uninformative metadata (e.g., “unknown”, “not applicable”, “not collected”, or blank entries) were excluded. Duplicate BioProjects and reference laboratory strains (e.g. *E. coli* K-12) were also removed to enrich for ecologically and clinically relevant diversity. Corresponding genome assemblies were downloaded from NCBI using the final list of accession numbers. All FASTA-formatted genome files were decompressed and concatenated into a single reference file to facilitate batch analysis. This curated set of genomes served as the foundation for all subsequent CRISPR array detection, sequence extraction, and conservation analysis.

### Identification of CRISPRs

CRISPR loci were identified across the filtered list of *E. coli* genomes using the standalone version of CRISPRCasFinder v4.2.20^41^. The tool was executed within a reproducible Apptainer (Singularity) container obtained from the CRISPR-Cas++ resource (https://crisprcas.i2bc.paris-saclay.fr/), ensuring consistent runtime environments across computing nodes. The input was a concatenated multi-FASTA file comprising all the genome assemblies. CRISPRCasFinder was run with default scoring parameters, and the inclusion of *cas* gene detection enabled (-cas flag). Arrays were filtered to include only those with at least three direct repeats and an evidence level of 4, indicating high-confidence predictions supported by strong repeat and spacer conservation, flanking sequences, and structural features. Output from CRISPRCasFinder included GFF3 annotation files, consensus repeat sequences, spacer counts, and full array structures for each genome. These results were parsed, and these arrays were used for downstream analysis.

### Alignment and phylogenetic visualizations of whole genomes and identified CRISPRs

Phylogenetic relationships among the genomes containing high-confidence CRISPR arrays were inferred using both whole-genome and CRISPR locus-specific sequence alignments. For whole-genome alignment, assemblies were processed using Parsnp v2.0.5^43^, with *E. coli* K-12 MG1655 (RefSeq accession: GCF_000005845.2) provided as the reference genome. All genome assemblies were placed in a single input directory, and the alignment was performed with verbose logging enabled. Core genome SNPs identified by Parsnp were used to construct a phylogenetic tree in Newick format. The resulting tree was annotated with sample-level metadata (organism, host, country, and year of isolation, with the number of CRISPRs identified and at high confidence) and visualized using Interactive Tree of Life (iTOL v6)^44^. Multi-locus sequence type (MLST) was determined by MLST typing at the Center for Genomic Epidemiology (http://www.genomicepidemiology.org/)^45^.

For CRISPR-specific analysis, all evidence level 4 CRISPR arrays identified by CRISPRCasFinder were extracted and concatenated into a multi-FASTA file. These sequences, consisting of alternating direct repeats and spacers (RSR units), were aligned using MAFFT v7.525^46^ with automatic strategy selection and parallel processing enabled. The resulting alignment was used to construct a maximum-likelihood phylogenetic tree with IQ-TREE v2.4.0^44^, using default model selection. CRISPR-based trees were visualized in iTOL with metadata alongside the whole-genome phylogeny for comparative interpretation.

### Repeat–Spacer–Repeat (RSR) unit generation and curation

Evidence level 4 CRISPR loci were parsed to generate Repeat**-**Spacer**-**Repeat (RSR) units from CRISPRCasFinder GFF3 annotations, using contig FASTA sequences when GFF attributes were insufficient. Loci annotated on the minus strand were reverse-complemented and ordered consistently prior to unit construction. RSR units were defined as adjacent triplets (R1**-**Spacer**-**R2) and required valid repeat**-**spacer adjacency within each locus. Spacer sequences were quality filtered by length (20 – 40 bp) and nucleotide complexity (Shannon entropy of mononucleotide frequencies > 1.5)^47^. For within-genome presence/absence analyses, identical RSR sequences were deduplicated per genome by exact sequence identity of the concatenated unit. RSR units were merged with sample metadata (host, country, year) using genome accession base names, including expanding metadata fields that contained multiple accessions per record. Composite grouping variables were constructed to support stratified analyses (Host_Country, Year_Host, Year_Country, Host_Country_Year). Additional exploratory summaries and visualizations of spacerome composition and motif structure are provided in the accompanying project repository and documented.

### Spacer sharing, similarity matrices, and network construction

Spacer sharing was quantified across metadata-defined groups using set-based overlap of deduplicated spacer sequences. For each grouping variable, spacers were treated as presence/absence features per group (duplicates within a group removed), and pairwise sharing between groups was summarized by (i) the number of shared spacers and (ii) Jaccard similarity^48^,

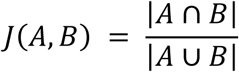

where 𝐴 and 𝐵 are spacer sets for two groups. Networks were constructed from the same pairwise overlap tables. Nodes represent metadata-defined groups; edges connect group pairs with at least one shared spacer and exceeding a specified Jaccard threshold. Networks were generated with Jaccard = 1.0 to capture more stringent conservation regimes. Edge weights were defined as Jaccard similarity. Community structure was inferred using Louvain modularity clustering on the weighted graph^49^, and node-level metrics (degree, betweenness) were computed; for weighted betweenness, edge “distance” was taken as the inverse of similarity 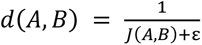 to reflect stronger similarity as shorter paths.

To distinguish non-random overlap from shared sparsity, edge-level significance was evaluated with a hypergeometric model over the universe of unique spacers observed in the analysis. For each pair of groups with spacer set sizes a and b, and intersection size k, enrichment was computed as

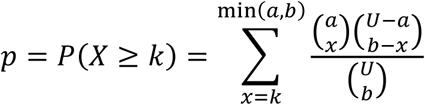

followed by Benjamini–Hochberg adjustment^47^. FDR-filtered networks were generated by retaining edges with BH-adjusted 𝑞 ≤ 0.05 in addition to the selected Jaccard threshold.

### Retrieval and geographic mapping of conserved spacerome of *E. coli*

To obtain additional *E. coli* genomes harboring conserved spacerome patterns, ten representative spacerome sequences determined in Tier 1 were selected. For each spacerome, the corresponding FASTA file of CRISPR spacerome was obtained from CRISPRFinder-derived outputs and used as the query set for BLAST searches. Searches were performed using the NCBI BLASTN tool with default parameters, except that percent identity was restricted to 100%, query coverage to 100%, and the E-value threshold set to 0 to ensure exact-matching spacerome sequences. All assemblies with complete matches to the ten query spaceromes were selected. For each matched genome, the WGS FASTA files and associated biosample metadata were downloaded from NCBI. The downloaded genomes then underwent CRISPR spacerome characterization as described above.

### Retrieval of WGSs for within STs

WGS of *E. coli* belonging to specific STs were retrieved from the NCBI Assembly database. A structured keyword-based query strategy was used to identify publicly available assemblies annotated with ST information. Searches were performed using organism- and ST-specific terms (e.g., “Escherichia coli”[Organism] AND “ST167”). For each ST, all searched assemblies were screened to confirm the presence of complete MLST metadata either in the Assembly record or in associated BioSample annotations. Only genomes with verified ST and “Complete Genome quality” were included. Corresponding FASTA files, GenBank annotations, and metadata were downloaded for subsequent CRISPR array extraction and comparative genomic analyses (**Table S3**). After profiling the CRISPR spaceromes, a spacer library was constructed to enable systematic and comparative analysis across strains (**Table S4**).

### Construction of the non-redundant protospacer database

A composite nucleotide database was constructed to trace *E. coli* spacer matches across diverse mobile genetic element and virome resources, including RefSeq viral, RefSeq plasmid, PLS-DB, Unified Human Gut Virome (UHGV), IMG/VR, and oral virome catalogs (HOVD, OPD, OED). Sequences from all sources were pooled and deduplicated to generate a non-redundant reference set, retaining a representative sequence for each exact-identity group and preserving a representative-to-member mapping to maintain provenance across sources. A BLAST+ nucleotide database was built from the non-redundant reference set, and source-specific metadata were harmonized into a unified master annotation table keyed to the reference identifiers.

### Proteospacer origin inference

Unique spacer sequences were queried against spacerome_NR using BLASTN configured for short sequences (blastn-short; -strand both, -ungapped, -dust no, -soft_masking false, -word_size 7, -max_hsps 1, -evalue 1e-5). Spacers were searched in length bins (20–22, 23–24, 25–27, 28–34, and ≥35 bp) and filtered to require full-length query coverage (100%) together with bin-specific minimum percent identity thresholds (90%, 93%, 95%, 97%, and 100%, respectively). For each spacer, the top hit was selected by descending bit score (ties resolved by longer alignment length). Hit subjects were annotated using the unified master metadata table, and representative-level annotations were propagated to member records using the representative-to-member mapping. To summarize origin structure across metadata-defined spacerome groups, hits were collapsed to an origin category based on reference provenance (viral, plasmid, other), and taxonomy was summarized at the class level using curated virus classifications when available and lineage-derived assignments otherwise. Alluvial (Sankey) diagrams were generated to visualize flows between spacerome group membership and inferred origin categories.

## Data availability

All the codes and data used were deposited in Zenodo. All the codes and detailed methodological explanations are available in the GitHub repository: https://github.com/biocoms/crispr_spacerome.

## Author Contribution

**Figure S1.**
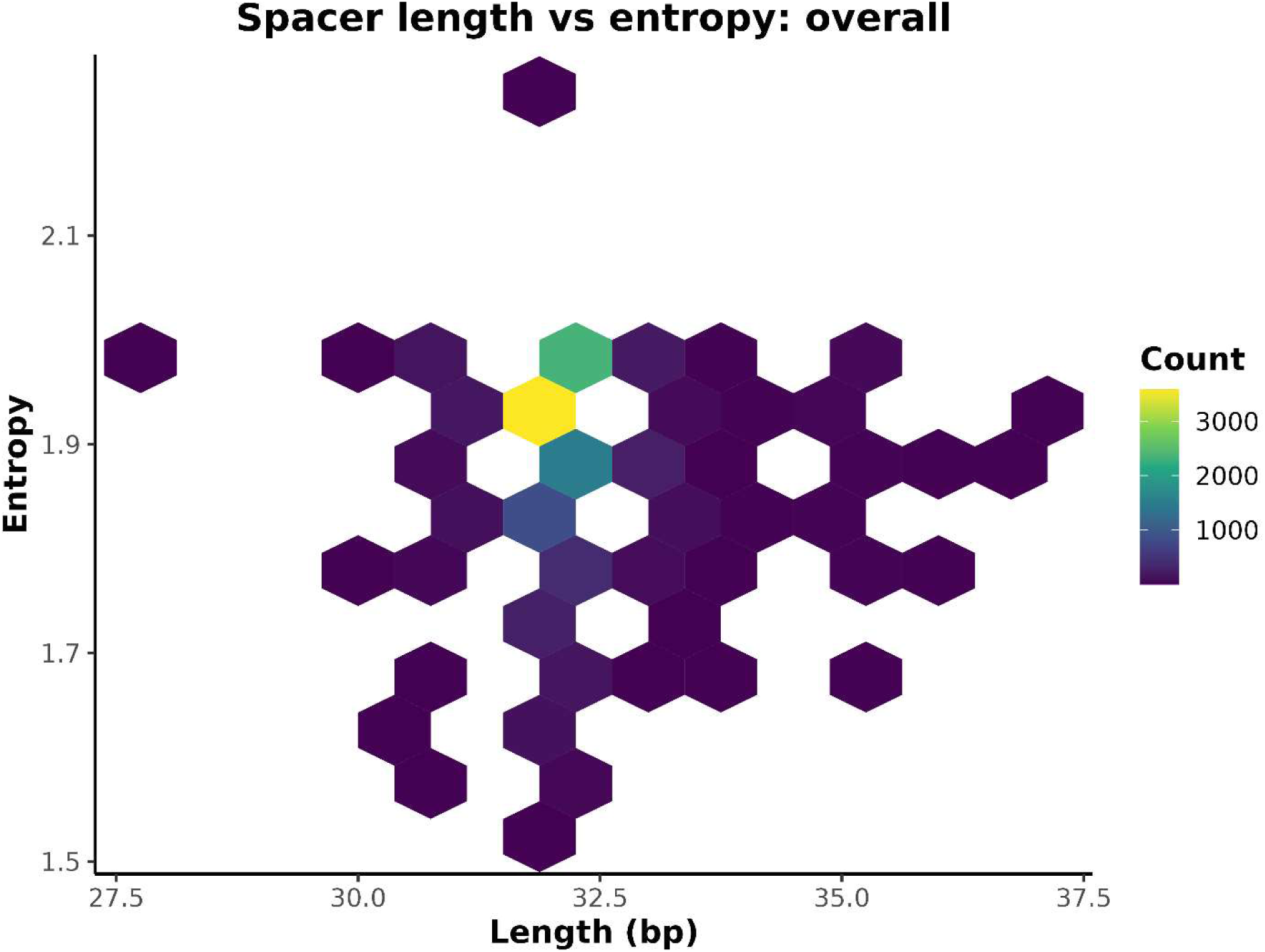
Spacer length versus sequence entropy across the *E. coli* CRISPR spacerome. Hexagonal binning summarizes the joint distribution of spacer length (bp) and Shannon entropy calculated from spacer nucleotide composition across all analyzed spacers. Color intensity reflects the number of spacers per bin, with warmer colors indicating higher counts.

**Figure S2.**
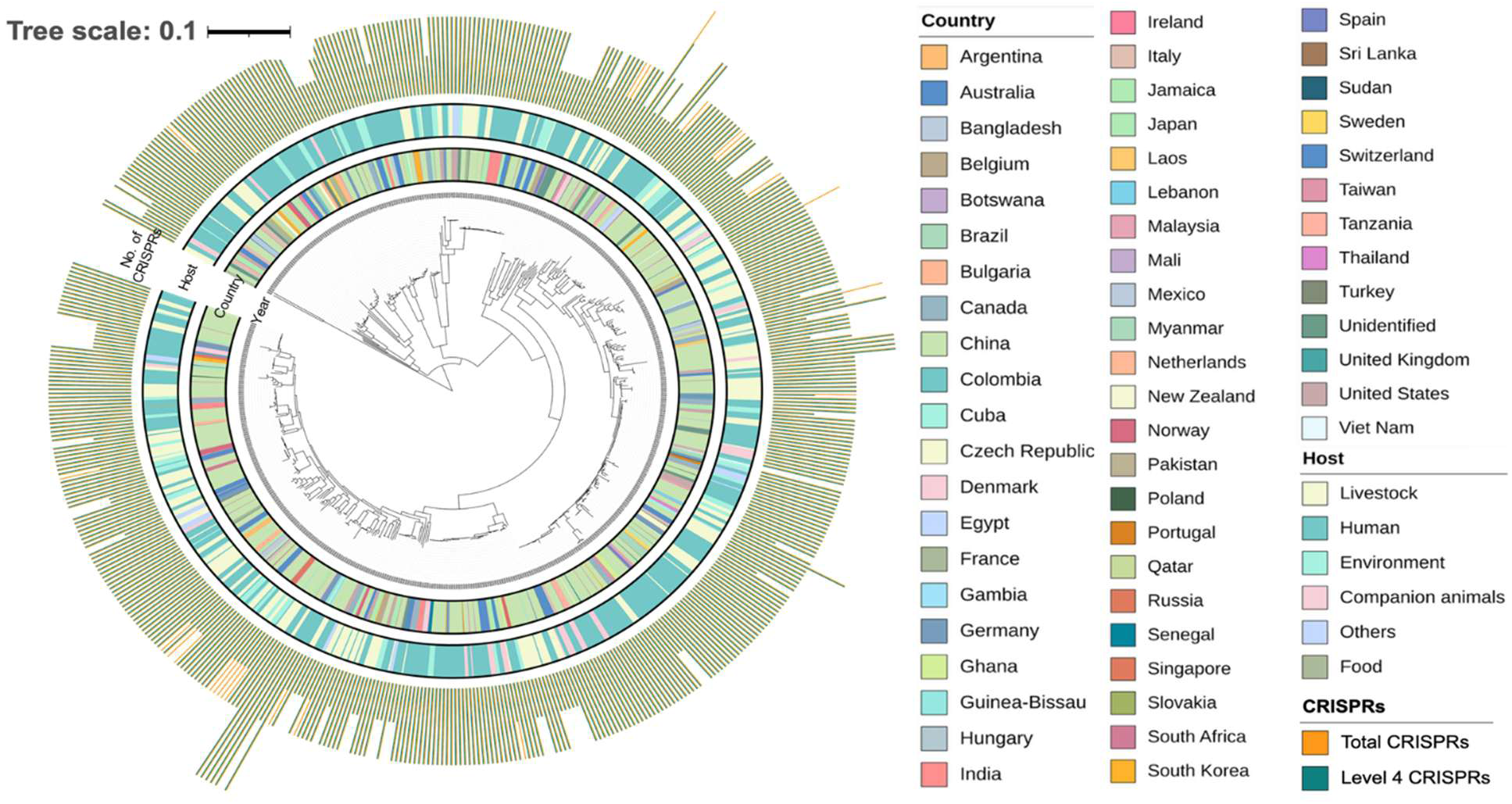
Global phylogenetic analysis of 532 *E. coli* genomes annotated with CRISPR spacerome features. The circular phylogenetic tree represents the core genome-based evolutionary relationships among 532 *E. coli* whole-genome sequences (WGS). The phylogeny was reconstructed using a SNP-based approach, and branch lengths are proportional to genetic relatedness. Each inner ring indicates the year and country of isolation. The middle ring denotes the host source, simply categorized as human, livestock, environment, companion animals, food, or others. The outer ring highlights the presence of CRISPRs. The outermost layer indicates the reliability of the identified CRISPR systems, with total identified CRISPRs shown in orange and high-confidence CRISPRs (Level 4) highlighted in teal.

**Figure S3.**
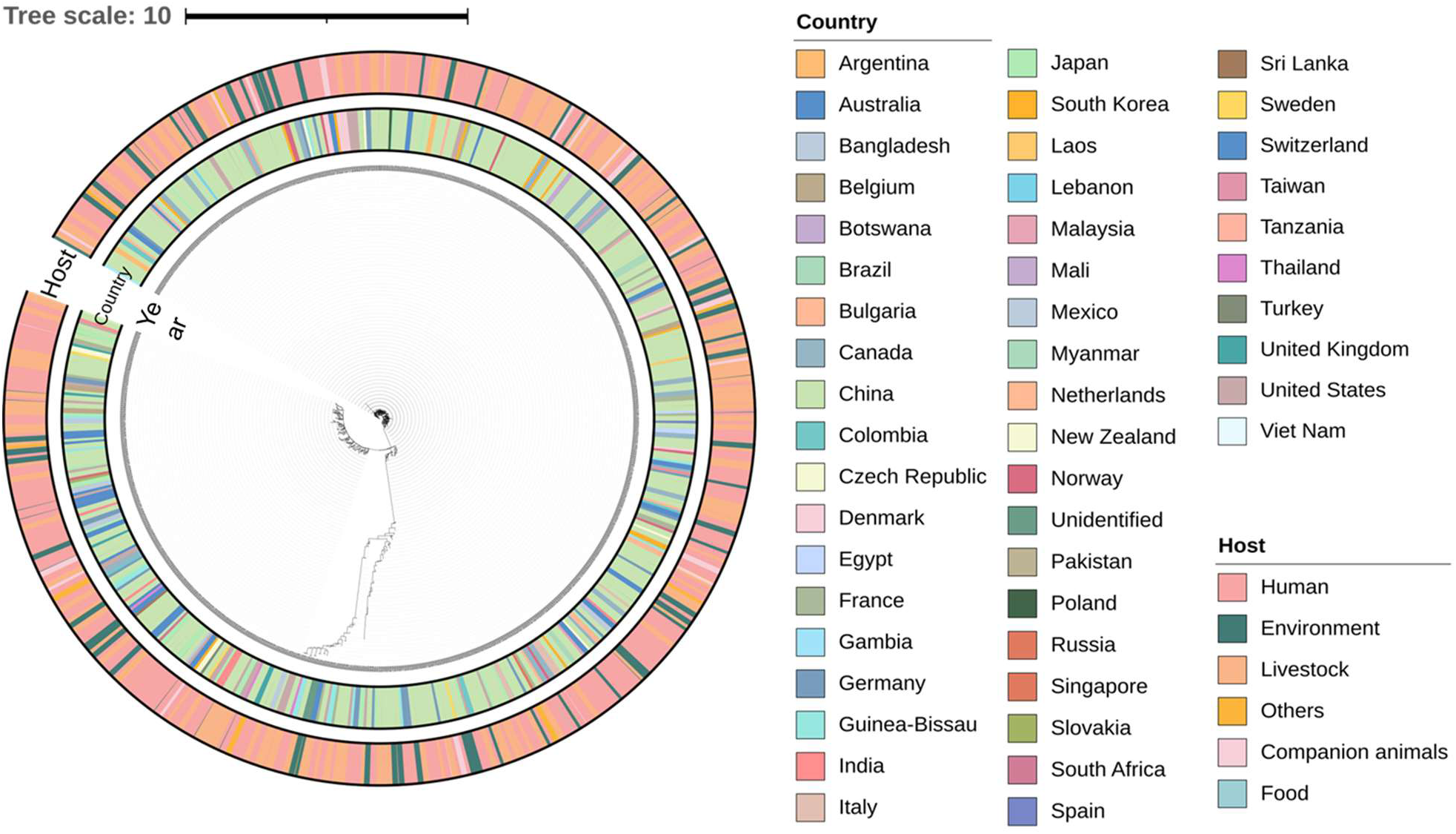
CRISPR spacerome-based genetic relationships among 532 global *E. coli* genomes. The circular phylogenetic tree represents relationships from 1,924 CRISPR spacerome sequences derived from 532 *E. coli* whole-genome sequences (WGS). The phylogeny was constructed based on spacerome sequence similarity, with branch lengths proportional to spacerome divergence. Inner and middle rings indicate the year and country of isolation. The outer ring denotes host source, categorized as human, livestock, environment, companion animals, food, or others. While this tree suggests the presence of closely related clusters, it does not confirm that the spacerome sequences are identical.

## Notes

### Competing Interest Statement

The authors have declared no competing interest.

### Funding

- Center for Health Effects of Environmental Contamination

## Reference

1. Hille, F. & Charpentier, E. CRISPR-Cas: biology, mechanisms and relevance. Philos. Trans. R. Soc. B Biol. Sci. 371, 20150496 (2016).

2. Makarova, K. S. et al. Evolution and classification of the CRISPR–Cas systems. Nat. Rev. Microbiol. 9, 467–477 (2011).

3. Shin, H. et al. Discovery of an identical CRISPR spacerome in a subset of global E. coli strains spanning decades. ISME Commun. ycaf211 (2025) doi:10.1093/ismeco/ycaf211.

4. Hanseob, S., Yongjin, K., Tatsuya, U. & Hor-Gil, H. Prevalence and Characterization of CRISPR Locus 2.1 Spacers in Escherichia coli Isolates Obtained from Feces of Animals and Humans. Microbiol. Spectr. 11, e04934–22 (2023).

5. Anderson, R. E. V & Boerlin, P. Carbapenemase-producing Enterobacteriaceae in animals and methodologies for their detection. Can. J. Vet. Res. = Rev. Can. Rech. Vet. 84, 3–17 (2020).

6. Shin, H. et al. Detection of a genetically related carbapenemase-producing Escherichia coli ST167 in clinical and environmental isolates: Evidence for clonal spread of carbapenemase-producing Enterobacteriaceae in humans and the environment in Iowa, United States. J. Glob. Antimicrob. Resist. 42, 154–160 (2025).

7. Petty, N. K. et al. Global dissemination of a multidrug resistant Escherichia coli clone. Proc. Natl. Acad. Sci. 111, 5694–5699 (2014).

8. R., M. A., et al. Global Extraintestinal Pathogenic Escherichia coli (ExPEC) Lineages. Clin. Microbiol. Rev. 32, 10.1128/cmr.00135-18 (2019).

9. Peirano, G. et al. Genomic Epidemiology of Global Carbapenemase-Producing Escherichia coli, 2015-2017. Emerg. Infect. Dis. 28, 924–931 (2022).

10. Watson, B. N. J. et al. CRISPR-Cas in Pseudomonas aeruginosa provides transient population-level immunity against high phage exposures. ISME J. 18, (2024).

11. Zhang, A.-N. et al. CRISPR-Cas spacer acquisition is a rare event in human gut microbiome. Cell Genomics 5, (2025).

12. Jee-Hwan, O. et al. Prophages in Lactobacillus reuteri Are Associated with Fitness Trade-Offs but Can Increase Competitiveness in the Gut Ecosystem. Appl. Environ. Microbiol. 86, e01922–19 (2019).

13. Muniesa, M., Colomer-Lluch, M. & Jofre, J. Potential Impact of Environmental Bacteriophages in Spreading Antibiotic Resistance Genes. Future Microbiol. 8, 739–751 (2013).

14. Burstein, D. et al. Major bacterial lineages are essentially devoid of CRISPR-Cas viral defence systems. Nat. Commun. 7, 10613 (2016).

15. Fehrenbach, A., Mitrofanov, A., Backofen, R. & Baumdicker, F. The complexity of multiple CRISPR arrays in strains with (co-occurring) CRISPR systems. bioRxiv 2025.05.05.651427 (2025) doi:10.1101/2025.05.05.651427.

16. Martynov, A., Severinov, K. & Ispolatov, I. Optimal number of spacers in CRISPR arrays. PLOS Comput. Biol. 13, e1005891 (2017).

17. Aydin, S. et al. Presence of Type I-F CRISPR/Cas systems is associated with antimicrobial susceptibility in Escherichia coli. J. Antimicrob. Chemother. 72, 2213–2218 (2017).

18. M., F. B., et al. Discovery of mcr-1-Mediated Colistin Resistance in a Highly Virulent Escherichia coli Lineage. mSphere 3, 10.1128/msphere.00486-18 (2018).

19. Laurence, C. M., J., R. C. & Philip, D. S. F Plasmid Lineages in Escherichia coli ST95: Implications for Host Range, Antibiotic Resistance, and Zoonoses. mSystems 7, e01212–21 (2022).

20. Roer, L., et al. Escherichia coli Sequence Type 410 Is Causing New International High-Risk Clones. mSphere 3, (2018).

21. D., P. J. D., Gisele, P., Yasufumi, M., Rebekah, D. & Liang, C. Escherichia coli sequence type 410 with carbapenemases: a paradigm shift within E. coli toward multidrug resistance. Antimicrob. Agents Chemother. 68, e01339–23 (2024).

22. Coque, T. M. et al. Dissemination of clonally related Escherichia coli strains expressing extended-spectrum beta-lactamase CTX-M-15. Emerg. Infect. Dis. 14, 195–200 (2008).

23. Christina, L. et al. O-Antigen Diversification Masks Identification of Highly Pathogenic Shiga Toxin-Producing Escherichia coli O104:H4-Like Strains. Microbiol. Spectr. 11, e00987–23 (2023).

24. Walker, L. L. et al. Emergence of a carbapenem-resistant atypical uropathogenic Escherichia coli clone as an increasing cause of urinary tract infection. Nat. Commun. 16, 8200 (2025).

25. Lopatina, A. & Sorek, R. CRISPR–Cas: Spacer Diversity Determines the Efficiency of Defense. Curr. Biol. 26, R683–R685 (2016).

26. Malina, A. et al. PAM multiplicity marks genomic target sites as inhibitory to CRISPR-Cas9 editing. Nat. Commun. 6, 10124 (2015).

27. Dion, M. B. et al. Escherichia coli CRISPR arrays from early life fecal samples preferentially target prophages. ISME J. 18, wrae005 (2024).

28. Song, S. et al. CRISPR-Cas Controls Cryptic Prophages. Int. J. Mol. Sci. 23, (2022).

29. Kim, M.-S. & Bae, J.-W. Lysogeny is prevalent and widely distributed in the murine gut microbiota. ISME J. 12, 1127–1141 (2018).

30. Kasman, L. M. Barriers to coliphage infection of commensal intestinal flora of laboratory mice. Virol. J. 2, 34 (2005).

31. Hansen, J., Gulati, A. & Sartor, R. B. The role of mucosal immunity and host genetics in defining intestinal commensal bacteria. Curr. Opin. Gastroenterol. 26, (2010).

32. Manson, J. M., Rauch, M. & Gilmore, M. S. The Commensal Microbiology of the Gastrointestinal Tract BT - GI Microbiota and Regulation of the Immune System. in (eds. Huffnagle, G. B. & Noverr, M. C.) 15–28 (Springer New York, New York, NY, 2008). doi:10.1007/978-0-387-09550-9_2.

33. Jiang, W. et al. Dealing with the Evolutionary Downside of CRISPR Immunity: Bacteria and Beneficial Plasmids. PLOS Genet. 9, e1003844 (2013).

34. Vale, P. F. et al. Costs of CRISPR-Cas-mediated resistance in Streptococcus thermophilus. Proc. R. Soc. B Biol. Sci. 282, 20151270 (2015).

35. Shkoporov, A. N., et al. The human gut virome is highly diverse, stable and individual-specific. bioRxiv 657528 (2019) doi:10.1101/657528.

36. Zorea, A. et al. Plasmids in the human gut reveal neutral dispersal and recombination that is overpowered by inflammatory diseases. Nat. Commun. 15, 3147 (2024).

37. Agarwal, A., Nayar, G. & Kaufman, J. Survey of Public Assay Data: Opportunities and Challenges to Understanding Antimicrobial Resistance. bioRxiv 2019.12.13.874909 (2019) doi:10.1101/2019.12.13.874909.

38. van Duin, D. & Doi, Y. The global epidemiology of carbapenemase-producing Enterobacteriaceae. Virulence 8, 460–469 (2017).

39. Johnson, J. R. et al. Global Distribution and Epidemiologic Associations of Escherichia coli Clonal Group A, 1998–2007. Emerg. Infect. Dis. J. 17, 2001 (2011).

40. Mitrofanov, A. et al. CRISPRidentify: identification of CRISPR arrays using machine learning approach. Nucleic Acids Res. 49, e20–e20 (2021).

41. Grissa, I., Vergnaud, G. & Pourcel, C. CRISPRFinder : a web tool to identify clustered regularly interspaced short palindromic repeats. 35, 52–57 (2007).

42. Collins, A. J. & Whitaker, R. J. CRISPR Comparison Toolkit: Rapid Identification, Visualization, and Analysis of CRISPR Array Diversity. Cris. J. 6, 386–400 (2023).

43. Kille, B. et al. Parsnp 2.0: scalable core-genome alignment for massive microbial datasets. Bioinformatics 40, btae311 (2024).

44. Letunic, I. & Bork, P. Interactive Tree of Life (iTOL) v6: recent updates to the phylogenetic tree display and annotation tool. Nucleic Acids Res. 52, W78–W82 (2024).

45. Larsen, M. V et al. Multilocus sequence typing of total-genome-sequenced bacteria. J. Clin. Microbiol. 50, 1355–1361 (2012).

46. Katoh, K., Misawa, K., Kuma, K. & Miyata, T. MAFFT: a novel method for rapid multiple sequence alignment based on fast Fourier transform. Nucleic Acids Res. 30, 3059–3066 (2002).

47. Haynes, W. Benjamini–Hochberg Method BT - Encyclopedia of Systems Biology. in (eds. Dubitzky, W., Wolkenhauer, O., Cho, K.-H. & Yokota, H.) 78 (Springer New York, New York, NY, 2013). doi:10.1007/978-1-4419-9863-7_1215.

48. Chung, N. C., Miasojedow, B., Startek, M. & Gambin, A. Jaccard/Tanimoto similarity test and estimation methods for biological presence-absence data. BMC Bioinformatics 20, 644 (2019).

49. Singlan, N., Abou Choucha, F. & Pasquier, C. A new Similarity Based Adapted Louvain Algorithm (SIMBA) for active module identification in p-value attributed biological networks. Sci. Rep. 15, 11360 (2025).

